# The RZZ complex integrates spindle checkpoint maintenance with dynamic expansion of unattached kinetochores

**DOI:** 10.1101/297580

**Authors:** Jose-Antonio Rodriguez-Rodriguez, Kara L. McKinley, Vitali Sikirzhytski, Jennifer Corona, John Maciejowski, Alexey Khodjakov, Iain M. Cheeseman, Prasad V. Jallepalli

**Affiliations:** Molecular Biology Program, Sloan Kettering Institute, Memorial Sloan Kettering Cancer Center, New York, NY, 10065, USA; Whitehead Institute for Biomedical Research, Nine Cambridge Center, Cambridge, MA 02142, USA; Department of Biology, Massachusetts Institute of Technology, Cambridge, MA 02142, USA; Wadsworth Center, New York State Department of Health, Albany, NY 12201; Rensselaer Polytechnic Institute, Troy, NY 12180

## Abstract

The Mad1-Mad2 heterodimer is the catalytic hub of the spindle assembly checkpoint (SAC), which controls mitosis through assembly of a multi-subunit anaphase inhibitor, the mitotic checkpoint complex (MCC) [1, 2]. Mad1-Mad2 first catalyzes MCC assembly at interphase nuclear pores [3], then migrates to kinetochores at nuclear envelope breakdown (NEBD) and resumes MCC assembly until bipolar spindle attachment is complete [1, 2]. There is significant debate about the factor(s) involved in targeting Mad1-Mad2 to kinetochores in higher eukaryotes [4-9]. Through gene editing and live-cell imaging, we found that the human Rod-Zw10-Zwilch (RZZ) complex is dispensable for cell viability and initial recruitment of Mad1-Mad2 to kinetochores at NEBD, but then becomes necessary to tether Mad1-Mad2 at kinetochores and sustain SAC arrest in cells challenged with spindle poisons. We also show that RZZ forms the mesh-like fibrous corona, a structural expansion of the outer kinetochore important for timely chromosome congression [10-13] once Mps1 phosphorylates the N-terminus of Rod. Artificially tethering Mad1-Mad2 to kinetochores enabled long-term mitotic arrest in the absence of RZZ. Conversely, blocking early RZZ-independent recruitment of Mad1-Mad2 eliminated the transient SAC response in RZZ-null cells. We conclude that RZZ drives structural changes in the outer kinetochore that facilitate chromosome bi-orientation and chronic SAC transduction, a key determinant of cytotoxicity during anti-mitotic drug therapy [14-16].

## Results

### RZZ is required for long-term mitotic arrest in response to spindle poisons

Recent studies have reached different conclusions about the specific roles and relative importance of Bub1 and the RZZ complex in targeting Mad1-Mad2 to kinetochores and transducing SAC arrest [4-8]. To interrogate RZZ function with maximum penetrance, we used AAV (adeno-associated virus)- and CRISPR-mediated gene editing to target the *KNTC1* (Rod) locus in HCT116 cells, a diploid colorectal cell line (Fig. S1A-C). The resulting *KNTC1*^HF/-^ (hypomorph-flox) cells expressed Rod at ~20% of the wildtype level (Fig. 1A-B and S1D) and exited mitosis prematurely when microtubule polymerization (nocodazole, 99 ± 6 min s.e.m.) or Eg5-dependent spindle bipolarity (S-trityl-L-cysteine (STLC), 193 ± 9 min s.e.m.) were inhibited, whereas wildtype cells never exited mitosis during the 16-hour timelapse (Fig. 1A-B). To determine if complete loss of the RZZ complex is compatible with clonogenic survival, *KNTC1*^HF/–^ cells were transduced with an adenovirus expressing Cre recombinase (AdCre) and subjected to limiting dilution.

**Figure 1.**
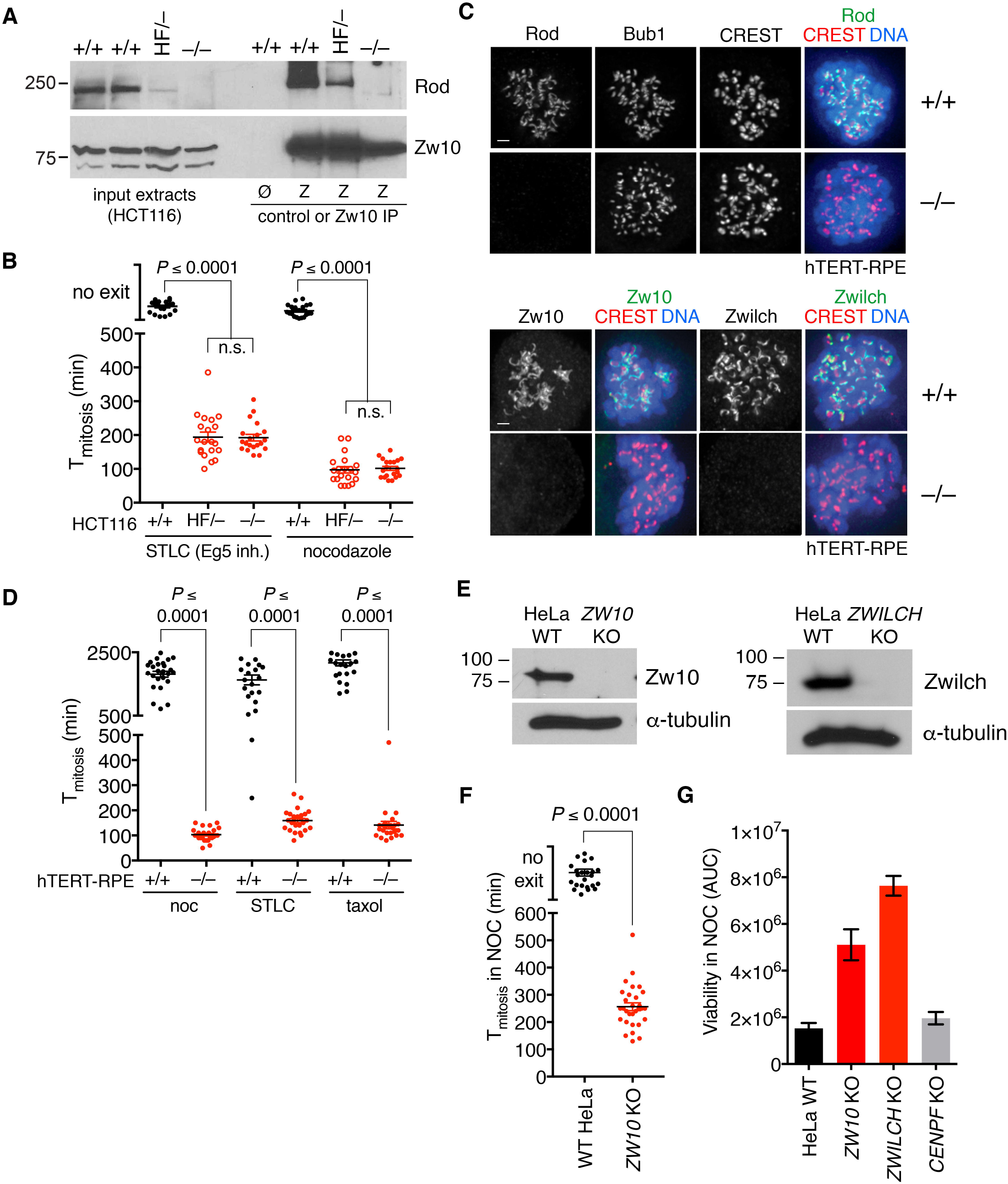
The RZZ complex is required for long-term mitotic arrest in response to spindle poisons. (A) AAV- and CRISPR-mediated genome editing was used to modify the *KNTC1* locus in HCT116 cells (see Fig. S1A-D). Cell extracts were prepared by nocodazole treatment, mitotic shakeoff, and nitrogen cavitation as described [3, 23, 34] and immunoprecipitated with Zw10-specific antibodies. Samples were resolved by SDS-PAGE and analyzed by Western blotting. (B) HCT116 cells expressing H2B-mCherry were treated with nocodazole or STLC and followed by dual-mode (epifluorescence and DIC) timelapse microscopy. Images were acquired at 10-min intervals. Mitotic duration (from NEBD to chromatin decondensation) was quantified in at least 25 cells per condition per experiment (N=2). P-values were computed using Kruskal-Wallis and Dunn’s multiple comparisons tests. Error bars indicate s.e.m. (C) Wildtype and *KNTC1*^−/−^ RPE1 cells (see Fig. S1E-F) were treated with nocodazole and MG132 for three hours, then fixed and immunostained to detect RZZ, Bub1, or human centromere autoantigens (CREST). Maximum-intensity projections of deconvolved z-series are shown. (D) *KNTC1*^−/−^ RPE1 cells were treated with nocodazole, STLC, or taxol and followed using DIC optics. Cell rounding (mitotic entry) and cortical blebbing and flattening (mitotic exit) were used as landmarks. (E and F) Clonal HeLa *ZW10* KO and *ZWILCH* KO cells were obtained from parental lines expressing gene-specific sgRNAs and doxycycline-inducible Cas9 [25]. Loss of Zw10 and Zwilch expression was confirmed by Western blotting. Mitotic duration was determined as in Fig. 1D. (G) Cells were treated with varying nocodazole concentrations (up to 660 nM) for 3 days and assayed for viability using CellTiter-Glo (Promega). Drug responses are expressed as area under the curve (AUC). *CENPF* KO cells were included as a control for Cas9 induction and cell cloning.

Unlike most SAC gene knockouts in mammals, *KNTC1*^−/−^ cells were viable (Fig. 1A-B). To extend these observations to other cell types, we deleted *KNTC1* in human retinal pigment epithelial (RPE) cells (Fig. 1C-D and Fig. S1E) and isolated *KNTC1*, *ZW10*, and *ZWILCH* KO clones from inducible knockout HeLa cells (Fig. 1E). Loss of any RZZ subunit abolished the kinetochore localization of its binding partners by immunofluorescence microscopy (IFM) and attenuated the SAC response to nocodazole, STLC, and taxol, which stabilizes microtubules (Fig. 1F and Fig. S1F-G). RZZ-deficient cells were also resistant to nocodazole-induced cytotoxicity (Fig. 1G), in line with previous studies correlating premature mitotic exit with resistance to taxanes and other antimitotic drugs [14-16].

### RZZ promotes chromosome congression and Mad1-Mad2 retention at unattached kinetochores

Basal mitotic timing was longer and more heterogeneous in RZZ-deficient cells (Fig. 2A), suggesting a delay in bipolar attachment and SAC satisfaction. Consistently, *KNTC1*-null cells rapidly exited mitosis when the SAC kinase Mps1 was inhibited with reversine (Fig. 2A). To monitor mitotic chromosome dynamics and SAC signaling, we expressed and imaged H2B-mCherry and FLAP-Mad1 using spinning disk confocal microscopy. At NEBD Mad1 immediately localized at nascent kinetochores, then dissociated as chromosomes congressed at the metaphase plate (Fig. 2B; n=10 cells). Congression was less efficient in *KNTC*-null cells, consistent with the lack of Spindly and dynein at kinetochores ([17] and Fig. S2), with small numbers of Mad1-positive chromosomes temporarily at the spindle poles (Fig. 2C; n=14 cells). We conclude that RZZ accelerates (but is not strictly required for) chromosome bi-orientation.

**Figure 2.**
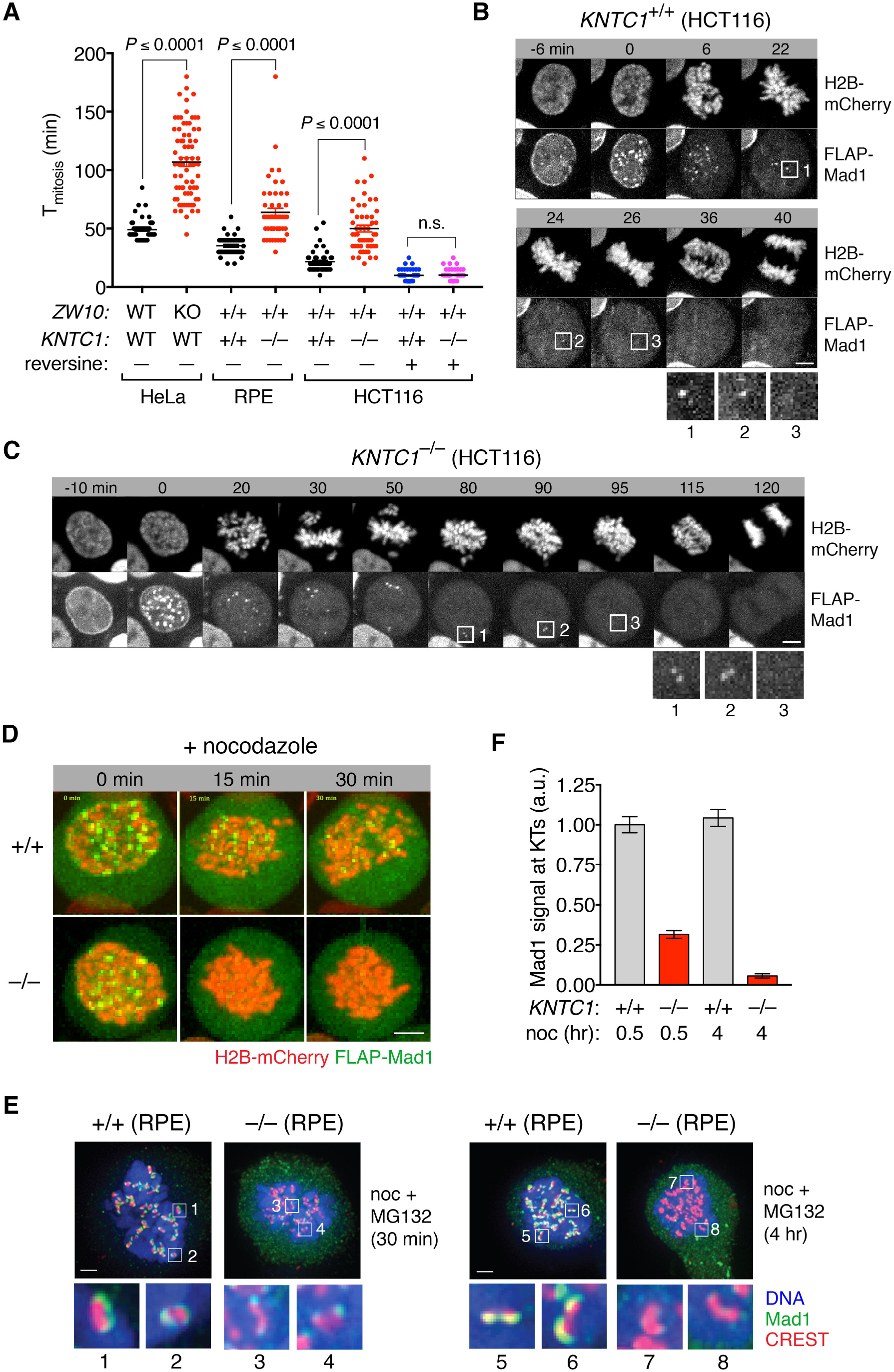
RZZ promotes chromosome congression and enables Mad1-Mad2 retention at unattached kinetochores. (A) Basal mitotic timing was determined for wildtype and RZZ-deficient HCT116, HeLa, and RPE1 cells using DIC timelapse microscopy. Where indicated, reversine was added to inhibit Mps1. Time 0 indicates NEBD. (B and C) Wildtype and *KNTC1*^−/−^ HCT116 cells expressing H2B-mCherry and FLAP-Mad1 were filmed during unperturbed mitosis using spinning disk confocal microscopy. Insets highlight coordination of chromosome congression, Mad1 dissociation from kinetochores, and anaphase onset in both cell lines. (D) Cells in (B and C) were filmed in the presence of nocodazole (n=6 for wildtype and n=14 for *KNTC1*^−/−^). (E and F) Wildtype and *KNTC1*-null RPE cells were treated with nocodazole and MG132 for 30 min or 4 hours before fixation for IFM. Insets in (E) highlight large Mad1 crescents in wildtype cells, versus small dim Mad1 foci in *KNTC1*^−/−^ cells. Mad1/CREST fluorescence intensity ratios were determined for at least 100 kinetochores in 5 cells per condition (N=3).

Next we examined Mad1 dynamics in nocodazole-treated cells. Although Mad1 was again targeted to kinetochores at NEBD, without RZZ its localization could not be maintained (Fig. 2D). To confirm this result for endogenous Mad1, wildtype and *KNTC1*^−/−^ RPE cells were treated with nocodazole and MG132 (to block mitotic exit) for 30 min or 4 hours, then fixed and analyzed by IFM. In wildtype cells Mad1 formed crescent-shaped structures that were stable over time (Fig. 2E, left panel). By contrast, in *KNTC1*-null cells Mad1 formed small dots that later disappeared from kinetochores (Fig. 2E, right panel and Fig. 2F).

### Mps1 phosphorylates the N-terminus of Rod to trigger kinetochore expansion

Mad1-Mad2 and RZZ localize to the fibrous corona, a phosphodependent expansion of the outermost kinetochore layer that persists until end-on microtubule attachment [12, 13]. Kinetochore expansion is thought to accelerate mitotic ‘search and capture’ by promoting lateral microtubule attachment [10] and to enhance SAC signaling [11]. RZZ is closely related to endomembrane coatomers that form oligomeric lattices [18, 19] and has been suggested as a “building block” of the corona itself [17, 20]. Consistently two other corona-associated proteins (CENP-E [21] and CENP-F [22]) also failed to form crescents in *KNTC1*^−/−^ cells (Fig. 3A). To obtain definitive evidence, we performed correlative light-electron microscopy in cells that also express CENP-A-GFP as a centromere marker. Serial sectioning revealed circumferential expansion of trilaminar plates and fibrous material in wildtype cells (n=14 kinetochores), whereas the kinetochores of *KNTC1*^−/−^ cells appeared as compact discs (n=15; Fig. 3B and Fig. S3).

**Figure 3.**
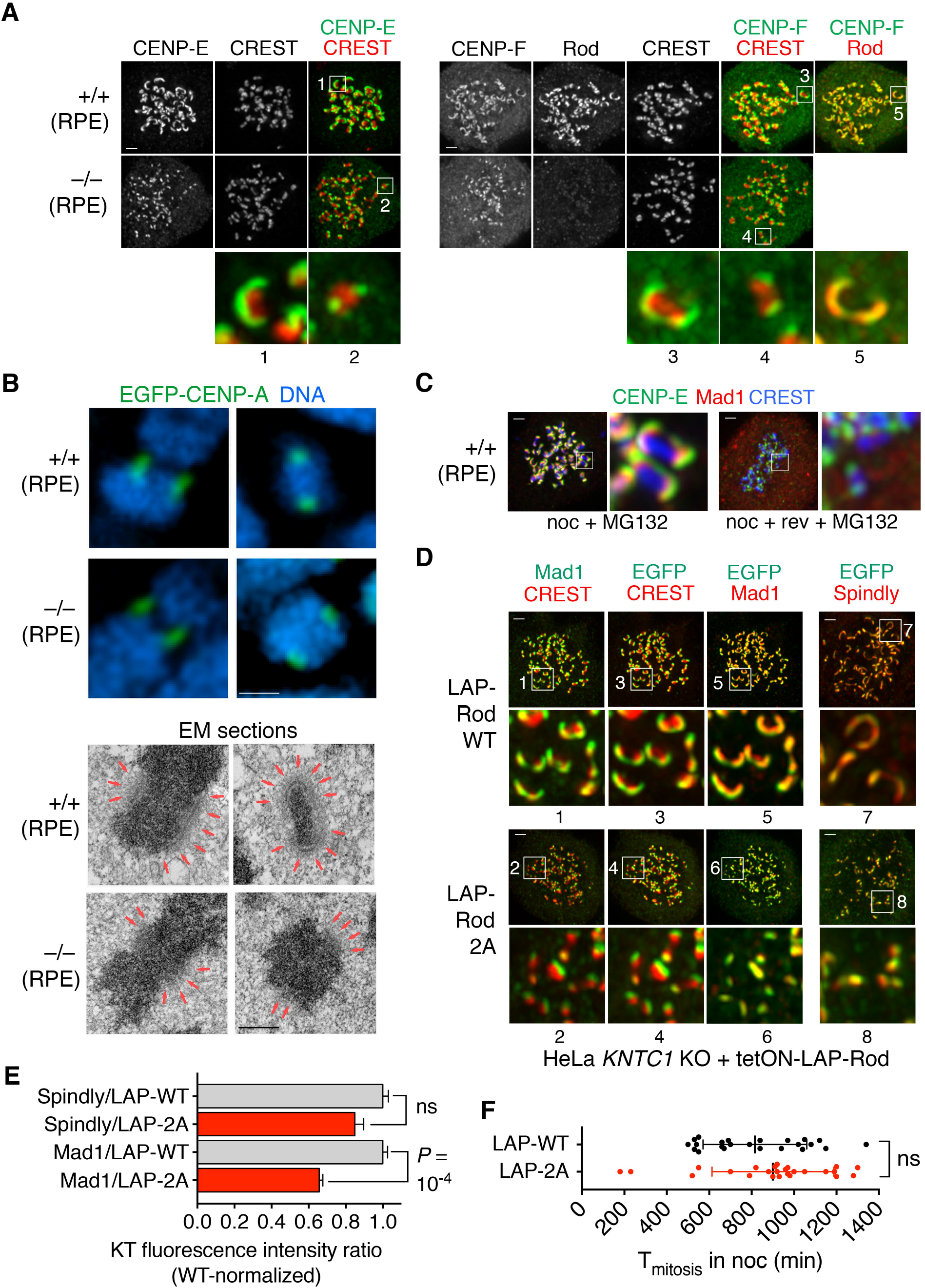
Mps1 triggers RZZ-dependent kinetochore expansion by phosphorylating the N-terminus of Rod. (A) Wildtype and *KNTC1*^−/−^ RPE cells were treated with nocodazole and MG132 for 2.5 hours before examining CENP-E, CENP-F, and Rod incorporation into crescents (n=5 to 10 cells, N=3). (B) Wildtype and *KNTC1*^−/−^ RPE cells expressing GFP-CENP-A were treated with nocodazole for 30 min and examined by correlative light and electron microscopy [53]. Upper and lower panels display LM and EM views of the same chromosomes. Red arrows mark the location and size of the outer kinetochore. Complete EM series are in Fig. S3. (C) RPE cells were treated with nocodazole and MG132 for 2 hr, after which reversine (rev) was added or omitted for 1 hr (n=10 cells, N=3). (D and E) *KNTC1*-null HeLa cells reconstituted with LAP-Rod^WT^ or LAP-Rod^2A^ were treated with nocodazole for 2.5 hr. IFM was used to visualize crescents and quantify MAD1/LAP-Rod and Spindly/LAP-Rod fluorescence intensity ratios (100 kinetochores in 5 cells each, N=2). (F) The duration of mitotic arrest in nocodazole-treated LAP-Rod^WT^ and LAP-Rod^2A^ cells was determined by DIC timelapse (n≥50 cells, N=2) and compared using the Mann-Whitney U test.

Kinetochore crescents disassembled when Mps1 was inhibited (Fig. 3C). We identified two Mps1 phosphorylation sites at the N-terminus of Rod (T13 and S15), upstream of its beta-propeller domain [23]. To test their function, wildtype (WT) and nonphosphorylatable (2A) Rod were expressed in T-Rex FLP-in HeLa cells as LAP (EGFP-TEV-S-peptide) fusions (Fig. S4A). Both versions were incorporated into RZZ based on co-immunoprecipitation assays (Fig. S4B). Thereafter we introduced a *KNTC1*-specific guide RNA (sg*KNTC1*) via lentiviral transduction and triggered gene disruption using a Cas9-expressing adenovirus (AdCas9). Endogenous Rod was removed from kinetochores in 75% of cells (Fig. S4C). We then analyzed transgene-expressing *KNTC1*-null clones. While LAP-Rod^WT^ and LAP-Rod^2A^ both localized to unattached kinetochores, only LAP-Rod^WT^ formed crescents (Fig. 3D). Thus far the only post-translational modification known to be required for crescent formation is C-terminal farnesylation of Spindly, which enables its kinetochore recruitment via interaction with Rod’s beta-propeller domain [17, 20, 24]. In contrast Rod^2A^ recruited Spindly normally (Fig. 3E). Despite recruiting less Mad1 to kinetochores, Rod^2A^ sustained mitotic arrest in nocodazole as effectively as Rod^WT^ (Fig. 3F). We conclude that Rod’s N-terminal phosphorylation activates Spindly-RZZ for kinetochore expansion but is not necessary for RZZ-mediated SAC transduction.

Bub1 promotes kinetochore expansion in *Xenopus* egg extracts [11] but has not been evaluated in human cells. We used doxycycline-inducible CRISPR [25] to delete *BUB1* in RPE cells with 60% efficiency as judged by IFM, with 42% of cells exiting nocodazole-induced mitotic arrest in less than 800 min (Fig. S4D-E). Large kinetochore crescents containing RZZ, Mad1, and CENP-E (but lacking CENP-F) formed in *BUB1*^−/-^ cells (Fig S4D). We conclude that Bub1 is not required for RZZ-mediated kinetochore expansion or Mad1-Mad2 recruitment, but is needed to localize CENP-F and potentially other corona-associated proteins.

### Tethering Mad1 to kinetochores allows long-term SAC maintenance without RZZ

SAC failure in RZZ-null cells correlates with Mad1-Mad2 loss from kinetochores (Fig. 2). However this did not exclude the possibility that RZZ has other essential roles in SAC signaling, such as activating Mad1-Mad2 for catalysis. To address this issue we expressed mCherry-Mis12-Mad1 [26, 27] from a doxycycline-inducible promoter in cells undergoing acute deletion of *KNTC1* (Fig. 4A), so that Mad1-Mad2 was tethered to kinetochores in an RZZ-independent manner (Fig. 4B). This caused a longer arrest in *KNTC1*-null cells (1138 ± 428 min) than wildtype cells (791 ± 285 min; Fig. 4C), potentially due to the lack of dynein-mediated stripping of SAC mediators from metaphase kinetochores [28-30]. To eliminate this effect and evaluate the SAC response to unattached kinetochores, we repeated these experiments in the presence of nocodazole and found no significant difference between *KNTC1*-null and wildtype cells (Fig. 4C). Thus, Mad1-Mad2 tethering is sufficient to explain how RZZ extends the timeframe of SAC surveillance and mitotic arrest by a factor of 10.

**Figure 4.**
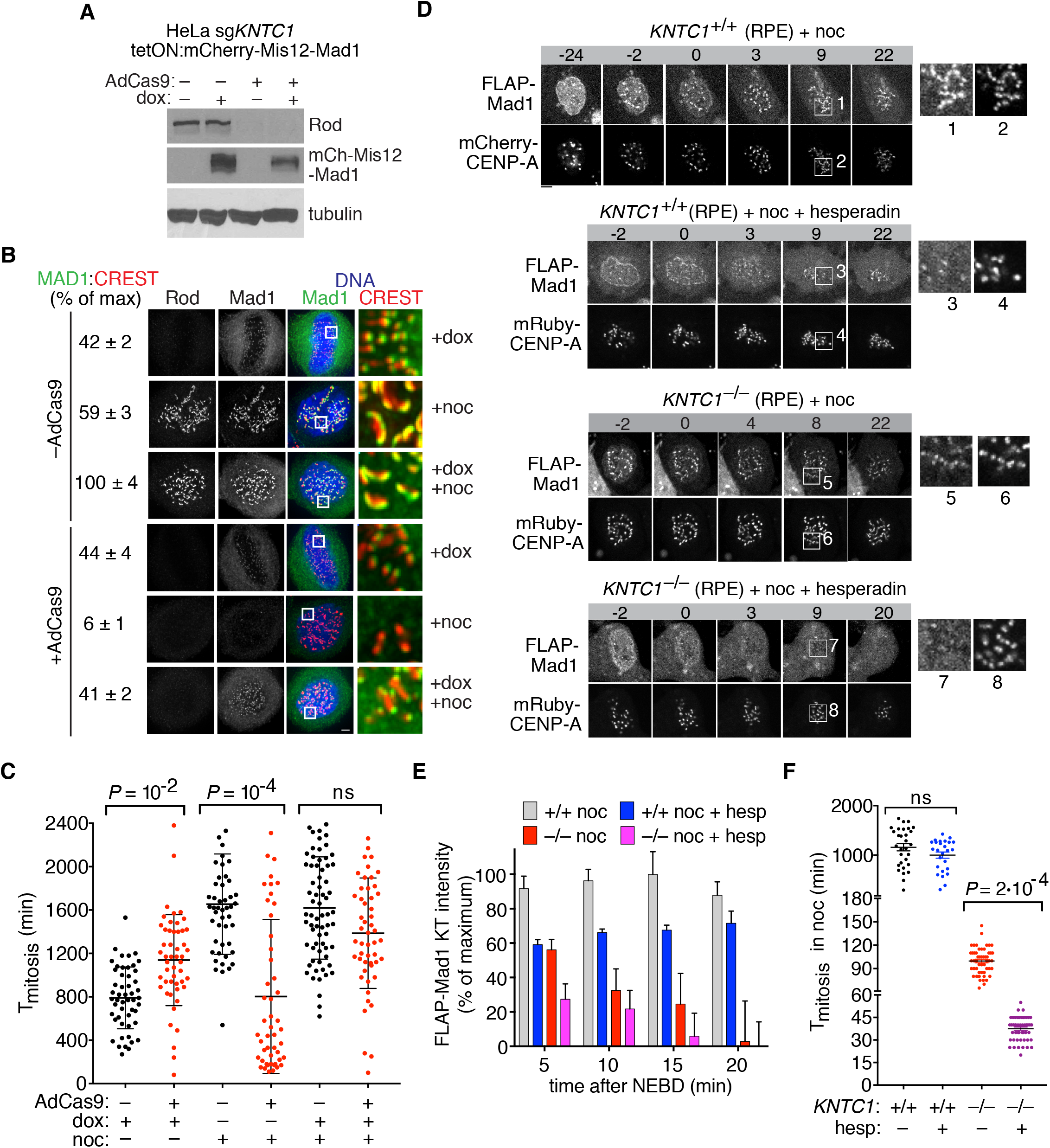
Aurora B and RZZ promote Mad1-Mad2 kinetochore recruitment and SAC transduction on complementary timescales. (A) HeLa cells expressing *KNTC1*-specific sgRNA and doxycycline-inducible mCherry-Mis12-Mad1 were infected with AdCas9 or mock infected. Three to five days later, cells were treated with or without doxycycline for 17 hr and analyzed by Western blotting. (B) Cells in (A) were analyzed by IFM, either with or without a further 3-hr treatment with nocodazole. Mad1/CREST ratios were determined for ≥100 kinetochores in 5 cells (N=2). (C) Cells in (A) were treated with doxycycline and/or nocodazole and followed by DIC timelapse imaging to quantify mitotic arrest (n≥50 cells, N=2). Error bars indicate s.d. (D and E) RPE cells expressing FLAP-Mad1 and mRuby-CENP-A were filmed during treatment with nocodazole ± hesperadin. Kinetochore-associated Mad1 was quantified from 5 timelapses per condition. (F) Mitotic arrest in nocodazole was quantified by DIC timelapse (n≥30 cells per condition).

### Aurora B and RZZ target Mad1-Mad2 to kinetochores and promote SAC transduction on complementary timescales

NPC-bound Mad1-Mad2 generates MCC during interphase, thereby increasing the SAC’s sensitivity and robustness [3]. This contribution was revealed when cells bearing an NPC-defective Mad1 allele were treated with Aurora B inhibitors, which delay kinetochore recruitment of Mad1 relative to NEBD [31, 32]. Under these conditions, cells entered mitosis without premade MCC and could not generate it quickly enough at kinetochores to prevent mitotic exit [3]. Because RZZ-deficient cells initiate but do not maintain the SAC, we anticipated a similar synthetic interaction with Aurora B inhibition. To test this idea, RPE cells co-expressing FLAP-Mad1 and mRuby-CENP-A were treated with nocodazole, either alone or in combination with hesperadin, an Aurora B-selective inhibitor [33]. Hesperadin slowed but did not prevent Mad1 recruitment to unattached kinetochores in wildtype cells (Fig. 4D-E) and did not override nocodazole-induced SAC arrest [3, 31, 33]. In contrast hesperadin blocked Mad1 targeting to kinetochores in *KNTC1*-null cells (Fig. 4D-E) and decreased mitotic arrest from 110 ± 20 min to 39 ± 10 min (Fig. 4F). As the latter corresponds to the basal length of mitosis in RPE cells (Fig. 2A), these data imply combined loss of SAC initiation and maintenance at kinetochores, while pre-programmed anaphase delay via nuclear pore-generated MCC remains intact.

## Discussion

The RZZ complex is conserved in metazoans and other taxa [18, 35, 36], but its roles in SAC signaling and kinetochore organization remain poorly understood. Our studies define RZZ as a critical SAC maintenance factor, the major Mad1-Mad2 tether at kinetochores, and a key mediator of phosphodependent kinetochore expansion.

Genetic loss of RZZ was tolerated in three different cell lines that vary in tumorigenicity, p53 proficiency, and karyotypic stability. In this regard RZZ differs from core SAC components like Mad1, Mad2, Bub1, BubR1, and Mps1 that are essential for organismal and (with rare exceptions [9, 37, 38]) somatic cell viability in mammals [3, 34, 39-42]. It was recently proposed that RZZ and Bub1 both transduce the SAC at unattached kinetochores, whereas only Bub1 monitors improper kinetochore-microtubule attachment [7]. However *KNTC1*-null RPE cells exhibited severe SAC failure when kinetochore attachments were ablated with nocodazole or when STLC or taxol were used to induce erroneous attachments (Fig. 1D). We conclude that RZZ is required for SAC transduction in both settings.

Through live imaging, we obtained evidence that human kinetochores recruit Mad1-Mad2 and transduce the SAC in two distinct steps. The first occurs at NEBD, when Mad1-Mad2 released from nuclear pores is targeted to newly assembled kinetochores by Aurora B [3, 31, 43]. In the second step, Mad1-Mad2 becomes stably tethered to kinetochores by RZZ, extending SAC surveillance and mitotic arrest more than 10-fold. When both steps are abrogated, cells neither initiate nor maintain the SAC and thus exit mitosis with kinetics set by nuclear pore-derived MCC [3].

Compelling evidence indicates that Bub1 is the kinetochore receptor for Mad1-Mad2 in yeast and nematodes [44, 45]. In contrast, studies in mammalian cells differ as to (1) whether Bub1 recruits Mad1-Mad2 to kinetochores and (2) whether and how Bub1 contributes to SAC arrest [4-7, 9, 41]. Recently several groups reported that the SAC in human cells is refractory not only to Bub1 depletion [4, 6] but also stable disruption of the *BUB1* locus [9, 46] unless Mps1 is partially inhibited to sensitize the system [9, 47]. In our studies, acute deletion of *BUB1* induced highly penetrant SAC failure without perturbing the kinetochore localization of RZZ and Mad1-Mad2 (Fig. S4D-E). These findings suggest that Bub1 supports SAC transduction by accelerating step(s) in MCC assembly downstream of Mad1-Mad2 [4]; however network-level compensation allows the SAC to adapt to chronic *BUB1* loss, similar to other signaling pathways [48, 49].

RZZ is also required for microtubule attachment-sensitive expansion of the outer kinetochore [10, 12, 13]. Transgenic expression of LAP-Rod^2A^ and mCherry-Mis12-Mad1 rescued nocodazole-induced SAC arrest but not kinetochore expansion in *KNTC1*-null cells, implying that the latter is not rate limiting for SAC transduction. This inference is supported by the fact that mitotic arrest following Eg5 inhibition (which generates compact kinetochores with syntelic attachments [50]) is also RZZ-dependent (Fig. 1B and 1D).

Although kinetochore expansion is known to depend on mitotic kinases like Mps1 [11], relevant substrates are not known. We found that Mps1 promotes expansion by phosphorylating Rod on two N-terminal residues upstream of its beta-propeller domain. Recent studies indicate that Rod and Spindly not only interact via this domain [17, 20] but also inhibit their own self-assembly into polymers [51, 52]. In light of these findings and the fact that Mps1 maximally phosphorylates Rod at unattached kinetochores [23], we propose that this modification overcomes the barrier to Spindly-RZZ polymerization and kinetochore expansion. Thereafter end-on microtubule attachments reverse this phosphorylation, in part through counteracting PP2A-B56 phosphatases [23], thus reinhibiting Spindly-RZZ and facilitating kinetochore compaction. In summary, we have identified a key trigger for dynamic kinetochore expansion that involves SAC-independent regulation of the RZZ complex by Mps1, further expanding the known substrates and functions of this mitotic kinase.

## Acknowledgments

We thank George Church, Didier Trono, David Sabatini, and Feng Zhang for sharing plasmids through Addgene. P.V.J. was supported by NIH grants R01GM094972 and P30CA008748.

## Author Contributions

J.-A.R.-R., K.L.M., J.C. and J.M. performed molecular biology, cell imaging, and biochemical studies and analyzed data. V.S. performed electron microscopy and analyzed data. A.K., I.M.C., and P.V.J. planned and supervised research, analyzed data, and secured funding. J.-A.R.-R. and P.V.J. wrote the paper with input from all authors.

## Declaration of Interests

The authors declare no competing interests.

### Supplemental Information – Movies

**Figure S1. AAV- and CRISPR-mediated deletion of *KNTC1* in human cells (related to Figure 1).** (A) Schematic of *KNTC1* locus, *AAV-KNTC1*^*HF*^ targeting vector, and wildtype allele-specific gRNAs used to delete exon 2 via CRISPR/Cas9. (B) Southern blot of PstI-digested genomic DNA hybridized to a [^32^P]-labeled 5’ probe (see panel A). (C) CRISPR-mediated deletion of exon 2 was verified by PCR. (D) Mitotic extracts were immunoprecipitated with Zw10-specific antibodies and immunoblotted as shown. (E) PCR assay confirming biallelic deletion of exon 2 in *KNTC1*^*−/−*^ RPE cells. (F) IFM confirms absence of Zw10 from kinetochores in HeLa *ZW10* and *ZWILCH* KO cells. (G) Taxol-treated HeLa *ZW10* and *ZWILCH* KO cells undergo premature mitotic exit.

**Figure S2. RZZ is required for kinetochore localization of Mad1-Mad2 and Spindly-dynein but not Bub1 or BubR1 (related to Figure 2).** Cells treated with nocodazole and MG132 for 2 hours (A-C) or untreated prometaphase cells (D-E) were analyzed by IFM as shown. Kinetochore-associated RZZ, Mad1-Mad2, Spindly-dynein (p150), Bub1, and BubR1 were quantified from ≥5 cells per condition and normalized against CREST (N=2).

**Figure S3. Serial EM sectioning of wildtype and RZZ-deficient kinetochores (related to Figure 3).** Full EM series for Fig. 3B. Red arrows indicate location and size of the outer kinetochore.

**Figure S4. Acute deletion of *KNTC1* and *BUB1* via inducible CRISPR.** (A) HeLa FLP-in T-Rex cells expressing a *KNTC1*-specific sgRNA and CRISPR-resistant LAP-Rod (WT or 2A) from a doxycycline-inducible promoter were infected with AdCas9 and/or induced with doxycycline as shown. Endogenous and LAP-tagged Rod were detected by Western blotting. Tubulin was used as a loading control. (B) Mitotic cell extracts from nocodazole-arrested HeLa cells expressing LAP Rod (WT or 2A) were immunoprecipitated with GFP-specific antibodies. Samples were resolved by SDS-PAGE and analyzed by Western blotting. Asterisk indicates a nonspecific cross-reacting band. (C) Cells in (A) were examined by IFM. Rod was not detected at kinetochores in 75% of AdCas9-transduced cells. (D) RPE iCas9 cells expressing a *BUB1*-specific sgRNA were induced with or without doxycycline for 3 days and treated with nocodazole for 2 hr. Kinetochore crescents were analyzed by IFM in 30 cells per condition (N=2). (E) Cells in (D) were filmed using DIC optics. Plot indicates cumulative mitotic exit in the presence of nocodazole.

Movie 1: Unperturbed mitosis in a wildtype HCT116 cell expressing FLAP-Mad1 and H2B-mCherry (related to Figure 2B).

Movie 2: Unperturbed mitosis in a *KNTC1*-null HCT116 cell expressing FLAP-Mad1 and H2B-mCherry (related to Figure 2C).

Movie 3: Mitotic entry in a wildtype HCT116 cell expressing FLAP-Mad1 and H2B-mCherry and treated with nocodazole (related to Figure 2D).

Movie 4: Mitotic entry in a *KNTC1*-null HCT116 cell expressing FLAP-Mad1 and H2B-mCherry treated with nocodazole (related to Figure 2D).

Movie 5: Mitotic entry in a wildtype RPE cell expressing FLAP-Mad1 and mRuby-CENP-A and treated with nocodazole (regulated to Figure 4D).

Movie 6: Mitotic entry in a wildtype RPE cell expressing FLAP-Mad1 and mRuby-CENP-A and treated with nocodazole and hesperadin (regulated to Figure 4D).

Movie 7: Mitotic entry in a *KNTC1*-null RPE cell expressing FLAP-Mad1 and mCherry and treated with nocodazole (related to Figure 4D).

Movie 8: Mitotic entry in a *KNTC1*-null RPE cell expressing FLAP-Mad1 and mCherry and treated with nocodazole and hesperadin (related to Figure 4D).

### Experimental Procedures

#### Cell lines and chemicals

Cell lines used in this study are described in the Key Resource Table. HeLa (human cervical adenocarcinoma, female) and HEK293 (human embryonic kidney, female) derivatives were grown at 37°C in Dulbecco’s modified eagle medium (DMEM) with 10% tetracycline free fetal bovine serum, 100 U/ml penicillin, and 100 U/ml streptomycin. hTERT-RPE (human retinal pigment epithelium, female) derivatives were grown at 37°C in a 1:1 mixture of DMEM and Ham’s F-12 medium with 10% fetal bovine serum, 100 U/ml penicillin, and 100 U/ml streptomycin, and 2.5 mM L-glutamine. Unless stated otherwise, nocodazole (660 nM), taxol (1 μM), S-trityl-L-cysteine (10 μM), MG132 (10 μM), hesperadin (100 nM), and reversine (500 nM) were used at the indicated concentrations.

#### Gene expression

LAP-Rod alleles (WT or 2A) were cloned into pcDNA5/FRT/TO. Constructs were cotransfected with pOG44 into HeLa T-Rex Flp-In cells using FuGene 6 (Roche). Integrants were selected using hygromycin (0.2 mg/ml), picked as single colonies, and induced with doxycycline (0.8 μg/ml). mCherry-Mis12-Mad1 [26] was cloned into a piggyBac vector containing a doxycycline-inducible promoter (*tetON*) and constitutively expressing reverse tetracycline transactivator (*rtTA*) and neomycin phosphotransferase (*neoR*) linked by the self-cleaving T2A peptide. HeLa cells were cotransfected with this construct and pSuperPiggyBac transposase (System Biosciences), selected in G418 (0.5 mg/ml), and induced as above. For stable expression of FLAP-Mad1, EGFP-CENP-A, and H2B-mCherry, retroviral transfer plasmids were cotransfected with pVSV-G into Phoenix 293 cells. For stable expression of mRuby-CENP-A or gene-specific sgRNAs, lentiviral transfer plasmids were cotransfected with psPAX2 and pMD2.G into Lenti-X 293T cells (Clontech). 24 to 48 hr later, supernatants were filtered, mixed 1:1 with fresh medium containing polybrene (20 μg/ml), and applied to target cells for 24 hr. Transductants were selected in G418, blasticidin (5 μg/ml), or puromycin (5 to 20 μg/ml).

#### AAV-mediated gene targeting

5’ and 3’ homology arms encompassing *KNTC1* exons 2 and 3 were amplified from human BAC clone RP11-18E11 using Pfusion DNA polymerase. A new *loxP* site was added upstream of exon 2 via XbaI digest and linker ligation. The entire targeting construct was transferred to pAAV as a NotI fragment. All manipulated regions were checked by sequencing to ensure their integrity. Procedures for preparing infectious AAV particles, transducing HCT116 cells, and isolating correctled targeted clones were performed as described [54]. In brief, Methods for AAV-mediated gene targeting have been described [54]. The *FRT-neo^R^-FRT* cassette was excised through transient expression of FLP recombinase (pCAGGS-FLPe) and limiting dilution. To delete *KNTC1*^HF^, cells were infected with AdCre (Vector Development Laboratory, Baylor College of Medicine).

#### CRISPR/Cas9-mediated genome editing

Zifit (http://zifit.partners.org/ZiFiT/ChoiceMenu.aspx) and sgRNA Designer (http://www.broadinstitute.org/rnai/public/analysis-tools/sgrna-design) were used to identify and rank candidate CRISPR/Cas9 targets for predicted on- and off-target activities. For transient expression, sequences were ordered as overlapping 60-nt oligonucleotides, annealed and extended into a 100-bp duplex using Pfusion DNA polymerase, and cloned into an AflII-digested gRNA expression vector (Addgene 41824) by Gibson assembly. Equal amounts of human codon-optimized Cas9 (Addgene 41815) and sgRNA vectors were transfected into HCT116 cells using FuGene 6 and into RPE cells using a Nucleofector 2b device (Lonza). For stable expression, target sequences were ordered as 24-nt oligonucleotides with asymmetric 5’ overhangs, phosphorylated with T4 polynucleotide kinase, then annealed and cloned into BsmBI-digested lentiGuide-puro (Addgene 52963) or pLenti-sgRNA (Addgene 71409) using T4 DNA ligase. Lentiviral transduction was performed as described above. Gene deletion was initiated by inducing a doxycycline-regulated Cas9 transgene present in the host cell line [25] or by infection with AdCas9 (ViraQuest).

#### Cell viability assays

Clonal HeLa *ZW10* KO and *ZWILCH* KO cell lines were isolated by limiting dilution after transient induction of Cas9 [25] and seeded at 1000 cells/well in black-walled 96-well plates. 3 days after adding nocodazole, cell viability was assessed using CellTiter-Glo (Promega) and quantified on a SpectraMax M5 microplate reader (Molecular Devices).

#### Immunofluorescence microscopy and live-cell imaging

Antibodies used in this study are listed in the Key Resource Table. Cells were fixed and permeabilized in PEMFT (20 mM PIPES, pH 6.8, 10 mM EGTA, 1 mM MgCl_2_, 4% paraformaldehyde, and 0.2% Triton X-100) for 13 min, blocked in 4% BSA, and stained with primary antibodies overnight. Species-specific secondary antibodies conjugated to Alexa Fluor 488, 564, or 647 were applied for 1 hr. Coverslips were mounted in ProLong Gold, imaged with a 100x oil objective on a DeltaVision Elite microscope (GE Life Sciences), and deconvolved in SoftWoRx using measured point spread functions.

For timelapse experiments cells were grown in multiwell plates or 35 mm glass-bottom dishes (MatTek) and imaged on a Nikon Eclipse Ti microscope equipped with a stage-top incubator and CO_2_ delivery system, 20x and 40x air objectives and 60x (1.4 NA) and 100x (1.45 NA) oil objectives, a Yokogawa CSU-X1 unit, and sCMOS (Andor Xyla 5.5) and EMCCD (Photometrics Evolve 512) cameras. Acquisition was performed with NIS Elements (v5.41). Epifluorescence and/or DIC images were acquired at 10-min intervals to measure mitotic arrest in response to spindle poisons, or at 2-min intervals to measure unperturbed mitotic timing. Confocal imaging of FLAP-Mad1, H2B-mCherry, and mRuby-CENP-A was performed at 2- to 5-min intervals in HCT116 cells (Fig. 2) and 1-min intervals in RPE cells (Fig. 4). Fluorescence intensities were quantified in ImageJ (v1.51) and analyzed in Prism 7.0 (GraphPad).

#### Correlative light-electron microscopy

Cells were fixed with 2.5% glutaraldehyde (Sigma) in PBS, pH 7.4-7.6 for 30 min, rinsed with PBS (3 × 5 min), and mounted in Rose chambers. Multimode (DIC and 3-color fluorescence) datasets were obtained on a Nikon TE2000 microscope equipped with a PlanApo 100x 1.45 NA objective lens at 53-nm XY pixels and 200-nm Z-steps. All LM images were deconvolved in SoftWoRx (v5.0) with lens-specific PSFs. Post-fixation, embedding, and sectioning were done as previously described [53]. Thin sections (70-80 nm) were imaged on a JEOL 1400 microscope operated at 80 kV using a side-mounted 4.0 megapixel XR401 sCMOS AMT camera (Advanced Microscopy Techniques Corp). Full series of images recorded at 10K magnification were used to reconstruct the volume of the cell, match orientation and superimpose this volume on the corresponding LM dataset. Higher-magnification images (40K) were then collected for individual kinetochores.

#### Cell lysis, immunoprecipitation, and Western blotting

Cell extracts were prepared by resuspending pellets in ice-cold buffer B (140 mM NaCl, 30 mM HEPES, pH 7.8, 5% glycerol, 10 mM sodium pyrophosphate, 5 mM sodium azide, 10 mM NaF, 10 mM PMSF, 0.3 mM sodium orthovanadate, 20 mM b-glycerophosphate, 1 mM DTT, 0.2 mM microcystin, and 1x protease inhibitor cocktail (Sigma)) prior to nitrogen cavitation (1250 psi, 45 min; Parr Instruments) and centrifugation at 20,000 x g for 30 minutes. LAP-Rod was immunoprecipitated with GFP antibodies coupled to protein G-Dynabeads using bis(sulfosuccinimidyl)suberate (BS3). Zw10 was immunoprecipitated without BS3 crosslinking. Extracts and immunoprecipitates were separated by SDS-PAGE and transferred to PVDF or nitrocellulose membranes. Membranes were blocked and probed with primary antibodies and secondary antibody-HRP conjugates in 5% nonfat dry milk in TBST (Tris-buffered saline + 0.05% Tween-20) before detecting signals via enhanced chemiluminescence (Western Lightning Plus, PerkinElmer).

